# Testing the maintenance of natural responses to survival-relevant calls in the conservation breeding population of a critically endangered corvid (*Corvus hawaiiensis*)

**DOI:** 10.1101/2021.05.24.445466

**Authors:** Anne C. Sabol, Alison L. Greggor, Bryce Masuda, Ronald R. Swaisgood

## Abstract

Vocal communication serves an important role in driving many animals’ social interactions and ultimately their survival. However, both the structure of and responses towards natural vocal behavior can be lost or subject to alteration under human care. Determining if animals in conservation breeding programs exhibit and respond appropriately to species-specific vocalizations is therefore important for ensuring their survival and persistence post-release. We tested whether endangered ‘alalā *(Corvus hawaiiensis),* which are extinct in nature, have retained their natural responses to vocal calls that were previously linked to survival and reproduction in the wild. We conducted our studies on breeding populations derived from a small number of founding ‘alalā maintained under human care since their extinction in the wild in 2002. We presented pairs of ‘alalā with alarm, territorial intrusion, and two types of control playback calls (a non-threatening territorial maintenance call and a novel heterospecific call). ‘Alalā were significantly more likely to approach the speaker following alarm call playback than other call types, and were more likely to respond to territorial intrusion calls with the same aggressive territorial calls. Males were more likely to make these aggressive calls than females, mirroring their roles in territory defense. We also found individual consistency in the level of vocal behavior response across all call types, indicating that some individuals are more vocal than others. These results are encouraging, showing that ‘alalā exhibit relevant, species-specific behaviors despite generations under human care. They do illustrate, however, that not all individuals respond appropriately, so animals’ responses to vocal stimuli may be an important factor to consider in determining the release suitability of individuals.

**Significance Statement:** Effective communication is crucial to the survival of many animals, but can erode without natural selection. Therefore, testing the flexibility and maintenance of communication and vocal responses in contexts where animals are isolated from conspecifics or from survival consequences, such as in conservation breeding centers, can help determine species’ susceptibility to communication loss. We used playbacks of survival-related conspecific calls to test if ‘alalā *(Corvus hawaiiensis),* retained species-specific responses to these calls after generations under human care. We found that birds maintained a species-level natural response, however these natural responses were not consistent across individuals, suggesting that some birds may not be well equipped to survive in the wild without additional training or care.

## Introduction

Animals have evolved mechanisms for communication that facilitate survival and reproduction. For primarily vocal species, how individuals respond to conspecific vocal cues and signals can have survival-relevant consequences. For instance, failing to respond to an alarm call could result in predation, or failing to adequately broadcast territorial calls could cause a loss of territory, mates, or offspring. Whether or not animals exhibit appropriate species-specific communication is particularly important in conservation breeding programs. Animals that have spent generations under human care need to retain their natural behaviors for successful return to the wild (Rabin 2003; McPhee 2004a, b). However, behavioral erosion is a common byproduct when animals are held under human care, resulting in the loss of survival-relevant behaviors, and deviations from wild-type behavior in form, or the expression of behaviors in inappropriate contexts. These alterations in behavioral phenotype can occur developmentally in a single generation or genetically/epigenetically across generations, and have been documented across a wide variety of vertebrate species for many functional categories of behavior, including antipredator, locomotory, foraging, refuge use, and reproductive, competitive and other social behaviors (Frankham 2008; Grueber et al. 2015, 2017). The loss of survival-relevant behaviors (McPhee and Carlstead 2010; Shier 2016), including losses in vocal diversity (Corfield et al. 2008; Digby et al. 2013) and dialect drift (Lewis et al. 2021) can alter call functionality and compromise reintroduction programs that use animals bred under human care. A combination of forces from evolution, culture (Brakes et al. 2019), and direct experience interact to shape animals’ vocal behaviors (Hollén and Radford 2009; Bradbury and Vehrencamp 2011), all of which may be impacted by the altered environment in conservation breeding facilities. Management of animals under conservation breeding, therefore, faces the challenge of providing opportunities for animals to maintain and express these behaviors in preparation for release into the wild (Greggor et al. 2018).

Among the vocal signals that animals need to retain, anti-predator signals (alarm calls) are particularly important because they are a component of an animal’s defense against predators (Hollen and Radford 2009). Developmentally, they may be more canalized, with alarm call production and responses emerging in development without direct experience with predators. While learning can be important for fine-tuning production and responses to antipredator calls (Seyfarth and Cheney 1980; Griffin et al. 2000), selection should favor individuals that respond appropriately to alarm calls without direct experience associating alarm calls with predation, therefore they should be relatively resistant to loss in comparison to other call types. However, some antipredator behavior responsiveness can be lost over an individual’s lifetime in the absence of predator exposure (Muralidhar et al. 2019). The importance of antipredator alarm calling for conservation breeding and translocation programs is underscored by the finding that predation is one of the primary causes of mortality for animals after release to the wild, across taxonomically diverse species (Fischer and Lindenmayer 2000; Moseby et al. 2011; Berger-Tal et al. 2020).

Another important set of signals helps animals avoid conspecific conflict by alerting others to shared or defended resources. Territorial species, for instance, have evolved multiple forms of communication that broadcast an intent to defend their territory. From ornamental displays, to physical demonstrations (e.g., Decourcy and Jenssen 1994), or auditory signals (Greenfield and Minckley 1993), there are a variety of ways that animals communicate territory occupancy to avoid unnecessary conflict. These calls often serve to communicate motivation and a level of severity of the territorial threat (Mager et al. 2012), ranging from maintenance calls to aggressive intrusion calls. The ability to efficiently set up and defend a territory can be critical to breeding and later reproductive success (Hiebert et al. 1989). Therefore, for territorial species that rely on vocal communication, having an ability to discern and respond to different territorial calls may be crucial for survival and reproduction in the wild. Any divergence in the production or perception of these important calls from wild phenotypes may challenge social integration and the success of translocation outcomes (Lewis et al. 2021).

Here we examine the vocal communication system of the endangered ‘alalā *(Corvus hawaiiensis)* within conservation breeding facilities, with the goal of evaluating their responses to major categories of conspecific vocal signals. Ultimately, we aim to determine if they have retained important components of natural behaviors that will facilitate successful reintroduction. The ‘alalā, or Hawaiian crow, is the last remaining corvid species of the Hawaiian islands (Banko et al. 2002). They were a keystone species for the Hawaiian wet and mesic forests, and the only known seed disperser for a number of native plants (Culliney et al. 2012). They went extinct in the wild in 2002, after decades of rapid decline due to disease, habitat loss/fragmentation and invasive predators, today surviving only in conservation breeding facilities (USFWS 2009). Efforts are ongoing to reintroduce ‘alalā back into the wild, yet the birds still face a long road to recovery and have not yet demonstrated successful breeding in the wild.

There have been but a few studies on ‘alalā vocal behavior, though we know they have a diverse vocal repertoire (Tanimoto et al. 2017a). Based on the complexity of vocal communication in other corvids (Enggist-Dueblin and Pfister 2002) and the role it plays in their social lives (Clayton and Emery 2007), we would expect ‘alalā calls to broadcast varying vocal signals, containing information about predators or conspecific territorial intrusions, with potentially important fitness consequences. ‘Alalā are fiercely territorial as adults, but as juveniles they form flocks, and associate with members of both sexes. Historically, breeding pairs, and especially males, would make a number of different, frequent territorial broadcast calls on the edges of their large territories (Banko et al. 2002); Additionally, like many species, ‘alalā are known to employ a range of alarm calls to warn others of danger (Tanimoto et al. 2017a; Greggor et al. 2021); although it is currently unknown if the structural differences between ‘alalā alarm call types are used to distinguish functional call categories. While comparatively little was known about ‘alalā calls when the species became extinct in the wild, evidence suggests that the frequency and type of some calls they make in the conservation breeding centers differ from their historical vocal behavior in the wild, including in the categories of territorial and alarm calls (Tanimoto et al. 2017b). It is unclear whether these changes are due to the erosion of natural behavior in the conservation breeding facilities, or due to a lack of context for expressing wildtype calls, for example, a lack of predation pressure resulting in a reduced need for alarm calls. Additionally, given that they are housed at much higher densities than wild ‘alalā and suffer reproductive consequences (Flanagan et al. 2020), it is possible that individuals in the conservation breeding population have become desensitized to their social surroundings, and no longer respond appropriately to these important signals, especially given that corvids have been shown to ignore unreliable callers (Wascher et al. 2015). Thus far there have been no studies examining behavioral responses to experimental playbacks of vocal calls in ‘alalā, beyond a preliminary pilot that identified alarm call types (i.e. Greggor et al. 2021). Given how little is known about the vocal behavior of ‘alalā in the wild before they went extinct, any future insight into call types and function must study the population under human care. We acknowledge that interpretations of the current fitness consequences of call function should be interpreted cautiously in light of these knowledge gaps.

We here conducted an investigation of ‘alalā responses to vocal signals to better understand vocal communication and behavior of this near-extinct species. Specifically, we examined whether ‘alalā still respond to species-specific calls in ways that indicate a retained meaning, despite their generations removed from nature, or whether their responses to conspecific call playbacks are indicative of a loss of vocal signals under human care, which might portend poor social integration and antipredator defense upon release. We presented birds with recorded playbacks of alarm and aggressive territory intrusion calls, alongside control calls and sounds, and measured how likely birds were to approach the calling sound and to respond to it with the same call type or a different call type (Table 1). We chose these two call types in the context of preparing birds for survival alongside predators, and the maintenance of social skills necessary for setting up and defending territories from conspecifics. We also presented ‘alalā with two control calls to test for a possibly confounding effect of the social novelty and auditory novelty of hearing a call outside of their aviary. Namely, we played a non-threatening ‘alalā territorial maintenance call, which birds routinely make from their existing territories, and should therefore not broadcast any threat, and a call from a novel species that ‘alalā have never heard.

**Table 1.** Explanation of stimuli types and predicted responses.

| Call type | Experimental design function | Predicted behavioral responses |
| --- | --- | --- |
| **Alarm** | Evaluate whether ‘alalā retain anti-predator responsiveness to conspecific calls indicating danger or predation threat | Vigilance, alarm calls, cautious approach (predicted to coordinate social anti-predator response) |
| **Territorial**<br>**Intrusion** | Discriminate between response to an alarming social stimulus versus an alarming antipredator stimulus | Aggressive approach, territorial intrusion calling |
| **Territorial**<br>**maintenance** | Control: non-alarming conspecific stimulus | No specific call response predicted, but ‘alalā could show interest and possible social attraction |
| **Novel Control** | Control: non-alarming heterospecific stimulus | No call response predicted; perhaps hesitancy to approach, based on neophobia |

If the environment of the conservation breeding facility has reduced birds’ responses to auditory stimuli generally (i.e. desensitization), we would expect that ‘alalā would produce no response to any of the call types (neither approaching nor making calls of their own). Responding naturally to one category, but not all of them, could indicate that some calls have lost their referential meaning either due to a lack of context for expression or as an artefact of generations of conservation breeding. Meanwhile, if birds have retained their natural responses to alarm and territorial intrusion calls, we would expect to see differences between them and control call types. Specifically, birds should respond to territorial intrusion calls with their own aggressive territorial intrusion calls (Bradbury and Vehrencamp 2011), and this effect should be most pronounced in males, due to the larger role males historically played in territorial defense (Banko et al. 2002). For alarm calls, predicting the natural response is more complicated. We would expect birds to approach the source of the call to investigate the potential source of danger, and perhaps to respond with their own alarm calls (Hill 1986; Manser et al. 2002). We predict birds should not show any clear response to the general territorial maintenance call, since it does not denote a threat, and may show signs of neophobia, or a hesitancy to approach, the novel call, since ‘alalā are highly neophobic in other contexts (Greggor et al. 2020).

## Methods

We conducted the experiment on 28 breeding pairs, i.e. 62% of the entire ‘alalā breeding population, at the Keauhou Bird Conservation Center (KBCC; n = 24 pairs) and the Maui Bird Conservation Center (MBCC; n = 4 pairs) between September and November 2018. The birds at both facilities are currently several generations removed from the wild. Birds were tested in their home enclosure with their breeding partner. Each pair was housed in an outdoor aviary, with covered areas for shelter and feeding, and had daily ad libitum access to food and water. Throughout the facility, interaction with people is minimized to reduce the effects of human care (see Greggor et al. 2018). The birds have auditory access to other breeding pairs but most do not have visual access. For the few buildings with more than one pair in adjacent aviaries we only tested one pair per building to minimize subjects’ prior exposure to the experimental setup. It is possible that birds could see the experimental setup at adjacent aviaries, so we put as much time as possible between trials at neighboring aviaries. At KBCC there was an average of 25 days between trials at neighboring aviaries, though due to scheduling constraints the range was somewhat large, between 5-53 days between trials at neighboring aviaries. At MBCC due to time constraints at this facility there were only two pairs of neighboring aviaries, and these trials were done 2 and 3 days apart. Four pairs were physically separated from each other for husbandry reasons, but both had visual access to each other during all of their trials.

### Experimental design of playback study

We designed the playback study to determine how ‘alalā respond to alarm and other social and non-social acoustic stimuli. We hypothesized that responses to different call types will vary according to the context in which they are used, and according to putative call meaning (Tanimoto et al. 2017a). The experiment entailed playback of four distinct acoustic stimuli: (1) conspecific alarm call, (2) conspecific territorial intrusion call, (3) conspecific non-threatening territorial maintenance call, and (4) heterospecific call from a novel bird species (Table 1). Alarm calls are high-pitched calls given to warn other birds of danger (Tanimoto et al. 2017a), which are typical of corvid vocal repertoires (e.g. Marzluff et al. 2010; Bila et al. 2016; Mclvor et al. 2018). By contrast, territorial intrusion calls are given when birds are actively and aggressively defending their territories, in situations that can escalate into physical aggression. Meanwhile, the non-threatening, territorial maintenance calls that ‘alalā make routinely on the edges of their territories, in the absence of any direct threat or aggression, were included as conspecific control stimuli.

We employed a repeated-measures design in which each subject received playback of each call type in a balanced, random order ensuring that each type of call was played first, second, third, or fourth an equal number of times across all trials. Individual and temporal differences in responsiveness to the experimental setup and acoustic playback can be a source of response variation that adds statistical noise in playback studies. To address this possibility, we included a pre-trial period before call playbacks in which subjects were exposed to ambient forest noise playback (details below). The same forest noise was used as a post-playback stimulus for all trials, so that conditions were constant for comparisons of observed behavior in pre- and post-call playback periods. We used this comparison to evaluate behavioral changes in the aftermath of call playbacks.

### Call recording and playback methods

We collected all audio recordings with a Roland R-05 acoustic recorder and directional boom microphone. Conspecific calls and forest noise controls were recorded at the KBCC. We collected calls opportunistically but tried to capture certain situations where we expected birds to make the calls in question. For example, for territorial intrusion calls we recorded when males were being moved between aviaries for husbandry purposes, as this is a time when we would expect these calls. Additionally, the alarm calls collected were verified against calls collected at times of recapture at the facility and during predatory exposure (e.g. Greggor et al. 2021). To minimize pseudoreplication, one set of calls was collected from each of three males, resulting in three unique sets of conspecific calls. A follow up analysis revealed no difference in birds’ responses between the three sets of calls (see Supplementary Materials). We assigned ‘alalā call types initially by social context and subjective perception, then confirmed assignment by examining spectrograms. Calls from unfamiliar species were collected opportunistically at the Panaewa Zoo in Hilo, Hawaii from medium- or large-bodied birds that breeding center ‘alalā would never have experienced (peacock *Pavo cristatus,* cockatoo *Cacatua moluccensis,* and toucan *Ramphastos toco).* Since several species of corvids are able to distinguish characteristics of conspecifics from calls (Boeckle et al. 2012; Mates et al. 2015; Woods et al. 2018), each pair received all three call exemplars from the same male. As breeding center birds are likely able to hear most conspecifics housed at the same facility, all subjects were likely somewhat familiar with individual calls from nearby conspecifics. Thus, to minimize familiarity with calls we selected playback calls from individuals that were not housed in adjacent aviaries and did not repeat call playback sets at adjacent aviaries.

We ensured that calls in the same category from different males had similar delta frequency and average power using Raven Pro, version 1.5. In order to standardize the duration and number of calls between call types with different durations and pauses, we ascertained that each 30-second stimulus contained 6-7 seconds of sound and between 6-20 individual calls, ensuring the natural spacing that the birds used between individual sounds to the extent possible. Therefore, we did not edit the calls themselves but did edit the spacing between calls and the number of calls to keep the total sound duration to 6-7 seconds for all playback stimuli. We measured sound duration and conducted all audio editing in Audacity. We also checked that all sounds in the same category were played from the speaker (omni jacket ultra, Altec Lansing) at the same minimum and maximum decibel level using the Decibel X app for iphones.

All playback trials were conducted between 9:30am and 11:30am, after morning husbandry-related disturbance had finished. This time window falls outside of the species-specific period of peak vocal activity that have been historically reported (0600 to 0900 and 1500 to 1800; Banko et al. 2002), and thus avoids times when background noise from other aviaries that might otherwise cause a distraction. For each playback trial we placed the speaker on the ground in a standardized location on the outside of the aviary, along an opaque wall. While the birds may have known a person was outside the aviary, they were unable to directly see the observer placing the speaker. We allowed the subjects a 5-min habituation period after the observer placed the speaker and moved to the observation location. Each of the four trials for each pair consisted of three different observation periods: pre-trial (3-min playback of forest noise to control for general response to sound playback), stimulus presentation (0.5-min call playback), and post-trial (3-min forest noise playback repeated). Following a 19-min intermission, we repeated the three-period trial using a different call type. This playback routine was repeated for all four calls. This resulted in 30 minutes between stimulus presentations. The observer scored trials live through a small window at the front of the aviary and recorded trials by setting up a video camera at each window. Video trials were consulted later to improve data accuracy. Because the stimuli were auditory and trials were scored live, it was not possible to record data blind. The birds likely heard the observer enter, but once the 5-minute habituation began, the observer stayed quiet and out of view as much as possible. Therefore, it is possible that the birds were aware of the presence of the observer and video camera, but we tried to minimize disturbance as much as possible. We recorded the following behaviors across all three trial periods: (1) approach (bird approaches playback speaker, measured as a binary variable with any movement of the bird in the direction of the speaker counting as an approach); (2) latency to approach (time at which bird first moved in the direction of the stimulus); (3) number and (4) type of all calls made by each bird. Call types were classified into four categories: alarm, territorial intrusion, subordinate begging and other. Territorial maintenance calls were never made in response to the stimuli so they were not included in our statistical models.

### Analysis

All analyses were conducted in R version 3.4.1 (R core team 2017). In order to determine if our pre-trial conditions were consistent for each stimulus, we ran a generalized linear mixed model (GLMM) with a negative binomial distribution (as data were zero inflated; Bliss et al. 1953) and a log link function, using the package glmmTMB (version 0.2.3; Brooks et al. 2017), with the number of all calls made during the pre-trial as the response variable. The initial model included the main effects of stimulus type and order as well as the interactions between stimulus type and order. We also included the random effect of bird ID.

To investigate the degree of individual consistency in call responses, we ran an intraclass correlation using the package irr (version 0.84; Gamer et al. 2012) for the number of calls each individual produced during the stimulus and post-trial combined for each stimulus type.

In order to determine whether birds responded differently to different stimuli types, we ran a cox proportional hazards model using the package survival (version 2.38; Therneau 2015) on the likeliness of birds to approach the stimulus for each stimulus type. This model included stimulus type, sex, and order as main effects as well as the interaction between stimulus type and sex and stimulus type and order. We clustered data around bird ID.

To evaluate the factors influencing call production, we used generalized linear mixed models (GLMM) with a negative binomial distribution (as data were zero inflated) and a log link function, using the package glmmTMB (version 0.2.3; Brooks et al. 2017) to separately test whether all calls, alarm calls, territorial intrusion calls and begging calls were more likely to occur during the stimulus and post-trial period depending on the playback type. All models initially included the main effects of stimulus type, sex, and trial order as well as the interactions between stimulus type and sex and stimulus type and order. We also included the random effect of bird ID. We used the territorial maintenance control call as the reference group when comparing the different stimulus types as this should represent a conspecific call in a new location, so this accounts for any calls or approaches simply due to the social novelty and not the call itself. The GLMM with the number of territorial intrusion calls made during the stimulus and post-trial would not converge properly when the model included interactions, so only main effects were tested.

For all GLMMs, with the exception of our model with territorial intrusion calls as the response variable, we first determined if the interaction terms warranted inclusion in the models. Starting with the interaction between stimulus type and order, as this was likely less biologically relevant, and then continuing with the interaction between stimulus type and sex, we removed interaction terms if their inclusion failed to decrease AIC values by > 2. We did not simplify the model past determining which interactions to include, as all remaining effects were important variables we wanted to consider in the final model. See the supplementary materials for the full process of model selection for each response variable. For all GLMMs we also visually inspected binned plots of the expected versus residual values for the final model.

## Results

### Pre-trial call frequency in response to playback of forest noise

As expected, we found no differences in call number among treatments before stimulus exposure during the pre-trial period (alarm trial vs. territorial maintenance: GLMM, b = −1.008, z = −0.942, *P* = 0.346; novel control trial vs. territorial maintenance: b = −0.836, z = −0.804, *P* = 0.421; territorial intrusion trial vs. territorial maintenance: b = −0.677, z = −0.665, *P* = 0.506). Therefore, we have not weighted post-trial data as a function of pre-trial calling rates. However, bird calls significantly increased across trials (b = 0.769, z = 2.179, *P* = 0.029); therefore, we included trial order in all subsequent models.

### Approach latency during stimuli presentation

Birds were more likely to approach after the alarm stimulus than the territorial maintenance stimulus (Fig. 1b; alarm vs. territorial maintenance: Cox Proportional Hazards Model, coefficient = 1.293, z = 2.293, *P* = 0.022). By contrast, the other stimuli had no significant effect on the likelihood of approaching relative to the territorial maintenance stimulus (Fig. 1a; novel control compared to territorial maintenance: coefficient = 0.827, z = 1.608, *P* = 0.108; territorial intrusion versus territorial maintenance: coefficient = 0.479, z = 0.892, *P* = 0.372). Trial order was also significant, with birds more likely to approach in later trials (coefficient = 0.242, z = 1.997, *P* = 0.0458). Sex did not have a significant effect on the likelihood of approaching (effect of sex: coefficient = 0.115, z = 0.376, *P* = 0.707).

**Fig. 1.**
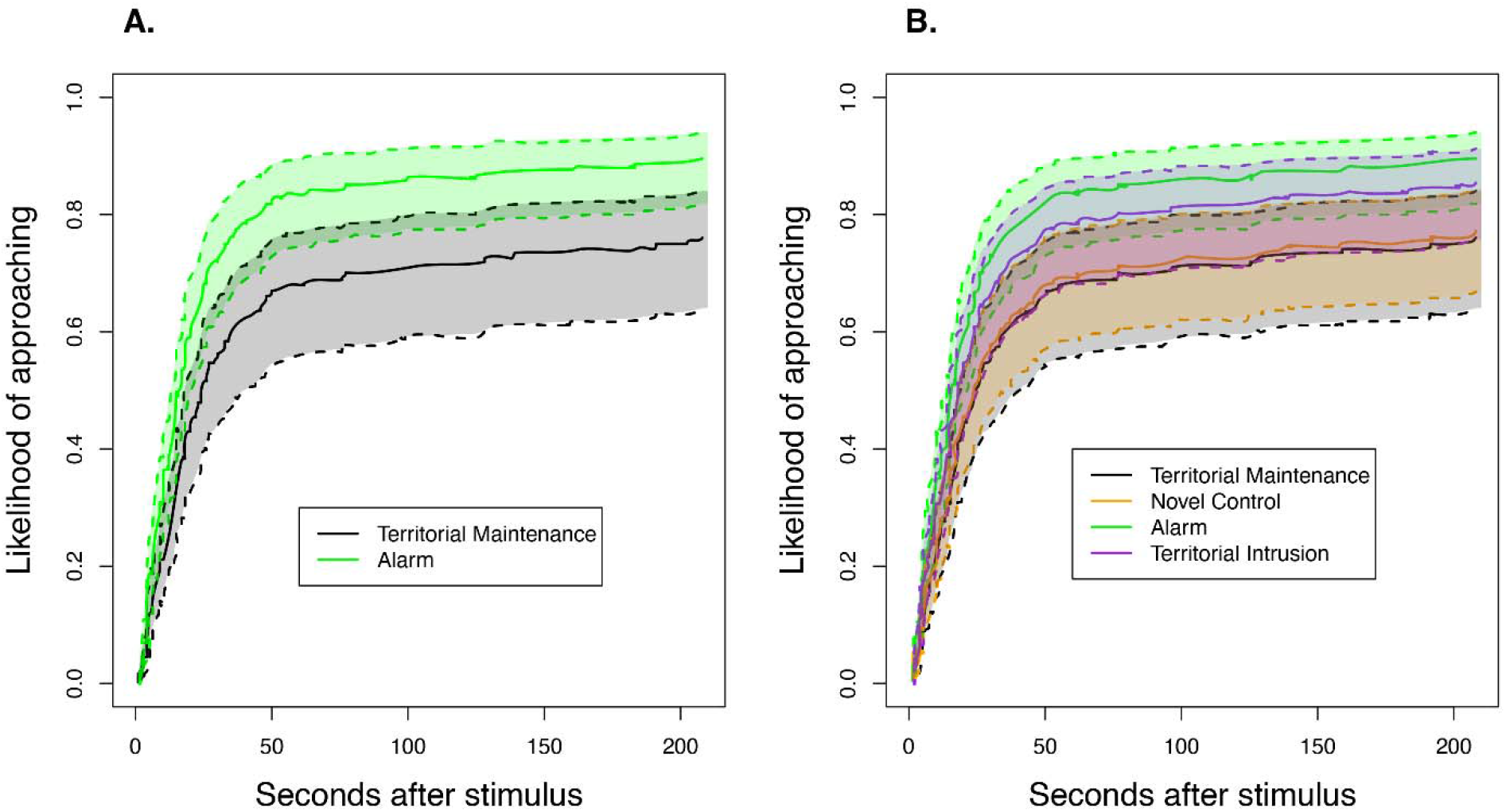
a Inverted survival curves (solid lines) showing the likelihood that birds approach in response to only the two significantly different stimuli, territorial maintenance and alarm (represented by the corresponding colors in the legend) with a 95% confidence interval (dotted lines). b Inverted survival curves (solid lines) showing the likelihood that birds approach each stimulus type (represented by the corresponding colors in the legend) with a 95% confidence interval (dotted lines)

### Individual call consistency during stimuli presentation and post-trial

Birds were individually consistent in the total number of calls they each made during the stimulus and post-trial periods across all different trial treatments (Intraclass correlation coefficient, 0.367, CI = 0.231-0.515, *P* < 0.001); i.e., birds who made fewer calls in one type of trial also made fewer calls in all trials.

### Call responses

Even though birds differed individually in the number of calls they made across all stimuli, no patterns emerged in how many calls birds made between stimuli types during the playback and post-trial period combined (Fig. 2; alarm vs. territorial maintenance: GLMM, b = 0.497, z = 0.737, *P* = 0.461; novel control vs. territorial maintenance: b = 0.271, z = 0.408, *P* = 0.683; territorial intrusion vs. territorial maintenance: b = 1.007, z = 1.471, *P* = 0.141). There was also no effect of sex (b = −0.598, z = −1.211, *P* = 0.226) or order (b = 0.099, z = 0.394, *P* = 0.693) on the total number of calls birds made during the stimulus and post-trial.

**Fig. 2.**
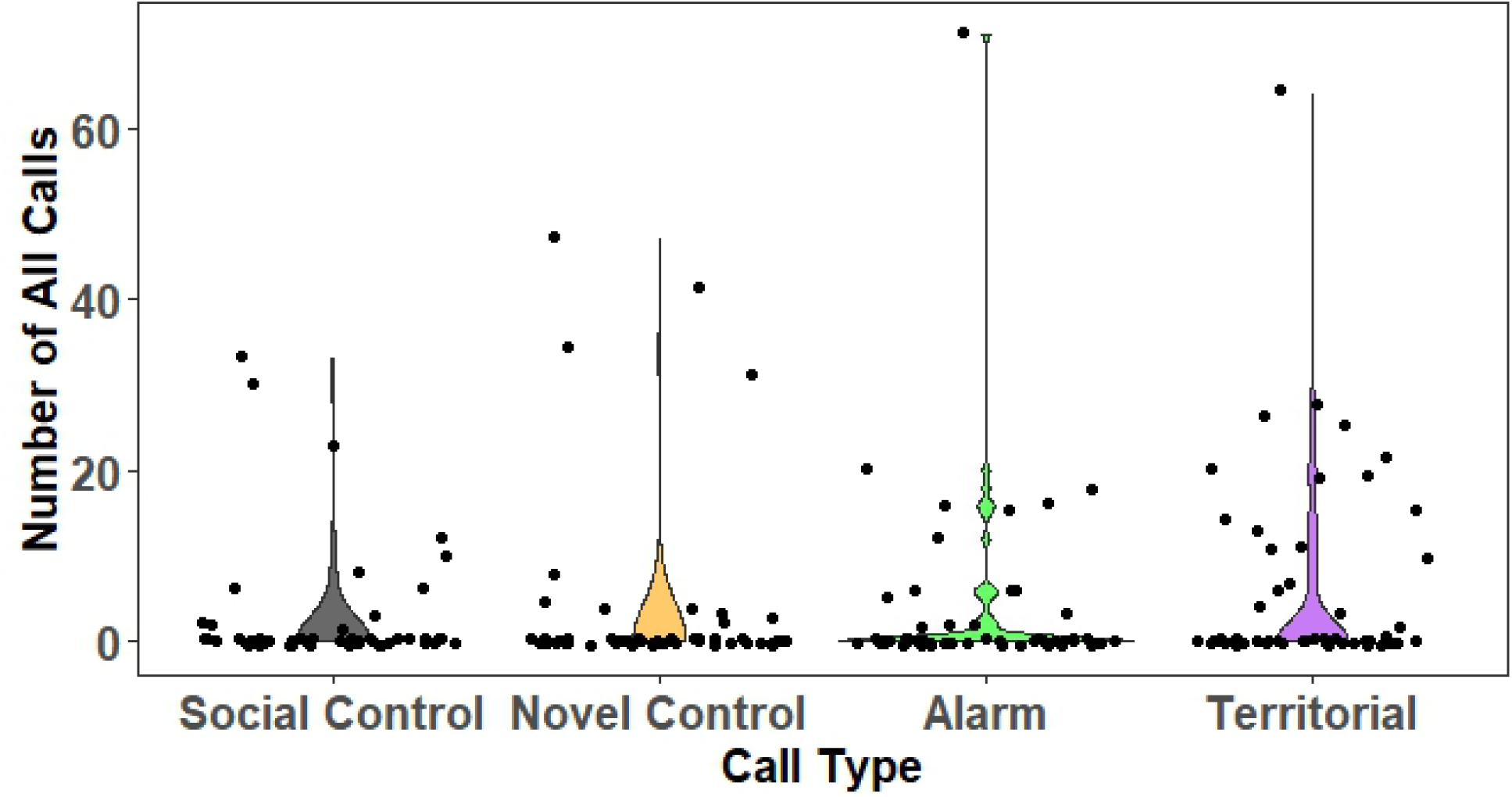
Violin plots and raw data showing the distribution of the number of all calls birds made during the stimulus and post-trial period after each type of stimulus. The points are the raw data for each individual trial, jittered to reduce point overlap

When we looked at the production of specific calls, we found that the number of alarm calls birds made during the stimulus and post-trial combined did not vary significantly between any of the different stimuli (Fig. 3; alarm stimulus vs. territorial maintenance: b = −1.584, z = −1.093, *P* = 0.275; novel control stimulus vs. territorial maintenance: b = 0.169, z = 0.097, *P* = 0.923; territorial stimulus vs. territorial maintenance: b = 0.390, z = 0.312, *P* = 0.755).

**Fig. 3.**
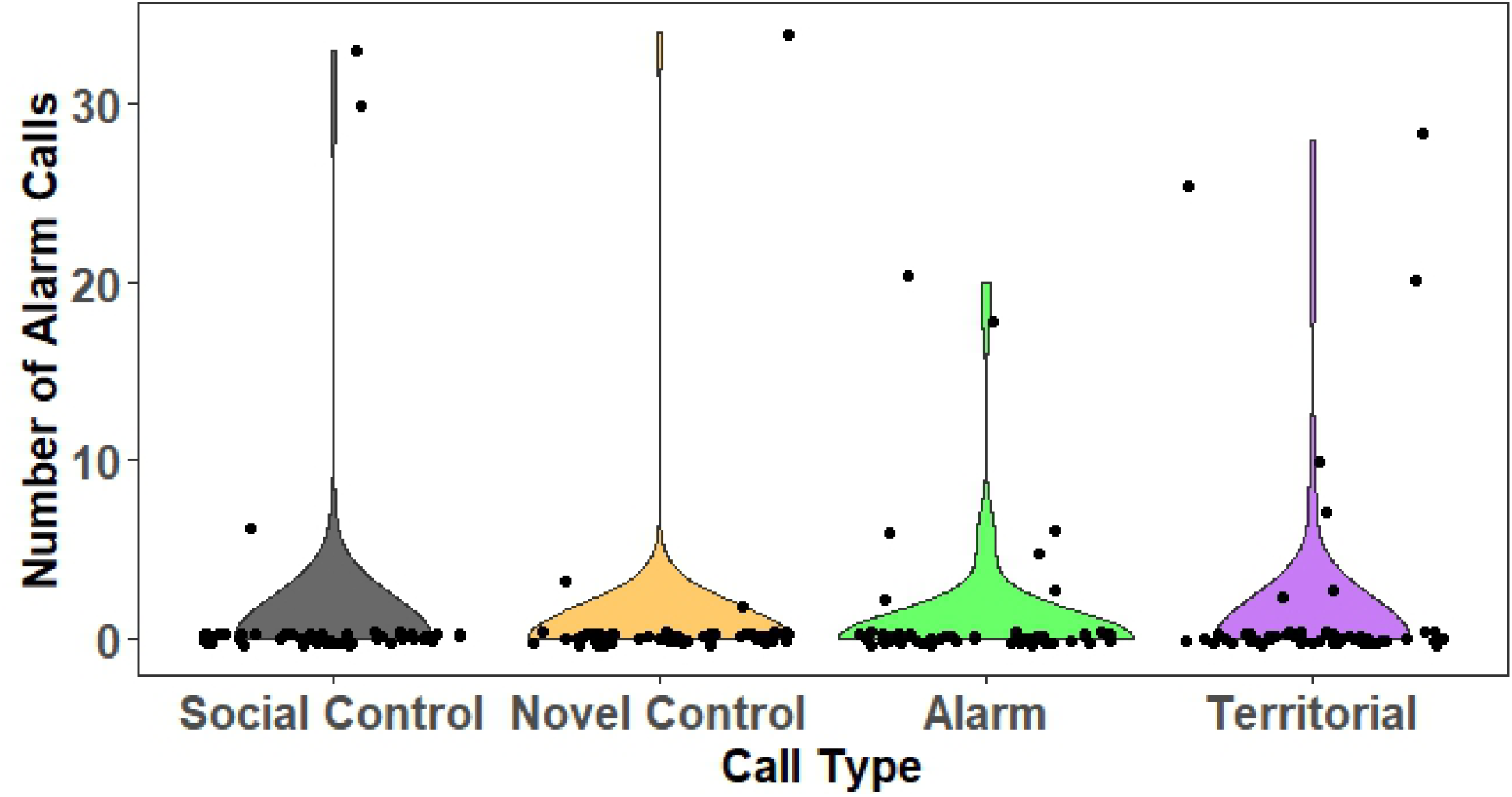
Violin plots and raw data showing the distribution of the number of alarm calls birds made during the stimulus and post-trial period after each type of stimulus. The points are the raw data for each individual trial, jittered to reduce point overlap

There was also no difference in the number of begging calls birds made during the stimulus and post-trial between any of the stimuli types (alarm stimulus vs. territorial maintenance: b = 0.820, z = 0.665, *P* = 0.506; novel control vs. territorial maintenance: b = −0.193, z = −0.170, *P* = 0.865; territorial intrusion stimulus vs. territorial maintenance: b = −0.441, z = −0.409, *P* = 0.683). Birds never made the territorial maintenance call in response to any stimuli.

Overall, birds made significantly more territorial intrusion calls during the stimulus and post-trial period in response to the territorial intrusion stimulus than to the territorial maintenance stimulus (Fig. 4; b = 3.825, z = 2.298, *P* = 0.022). By contrast, they did not make a significantly different amount of territorial calls during the stimulus and post-trial period in response to any other kind of stimulus (Fig. 4; alarm stimulus vs. territorial maintenance: b = 2.033, z = 1.064, *P* = 0.287; novel control vs. territorial maintenance: b = −19.797, z = −0.002, *P* = 0.999).

**Fig. 4.**
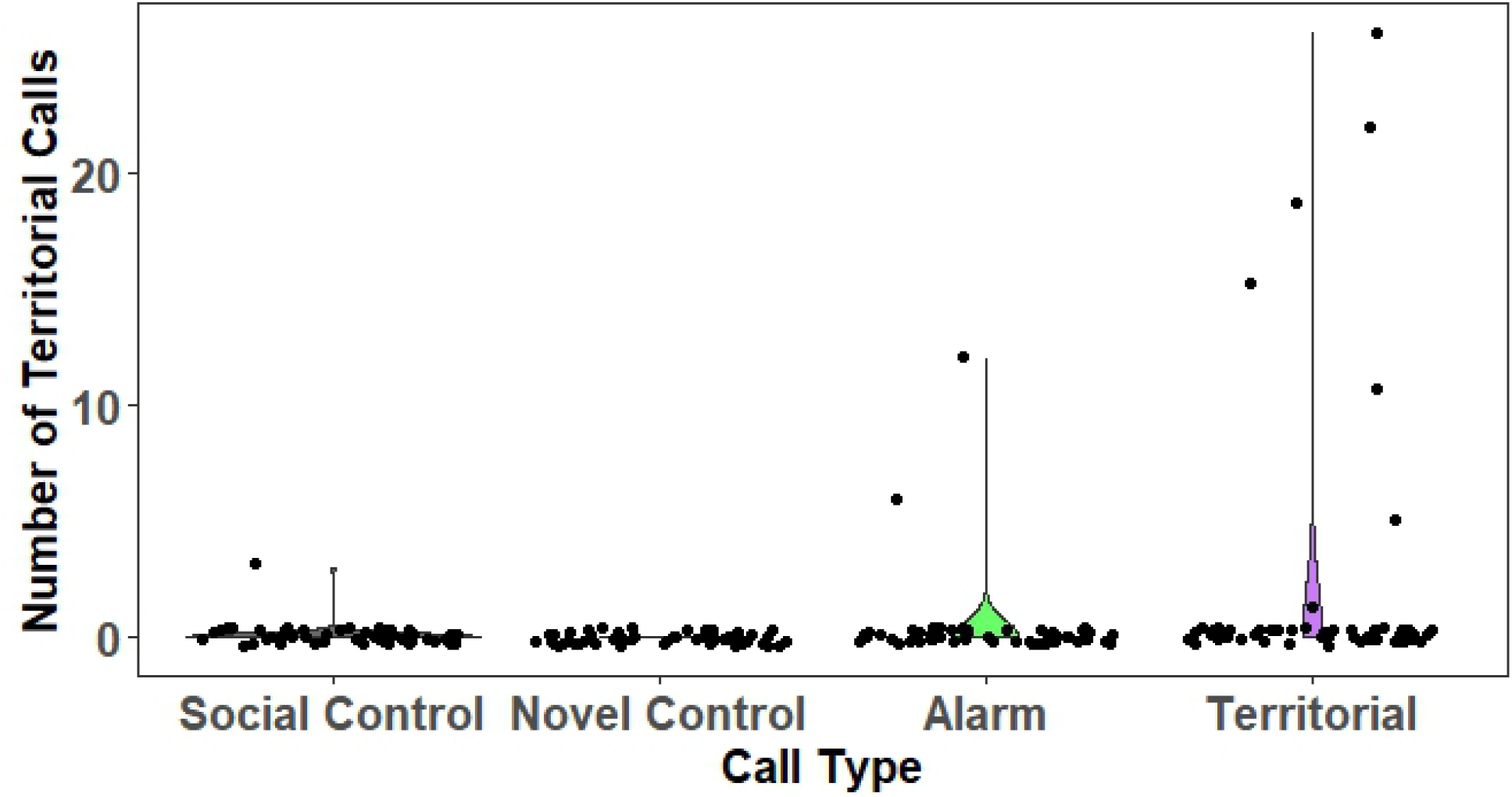
Violin plots and raw data showing the distribution of the number of territorial intrusion calls birds made during the stimulus and post-trial period after each type of stimulus. The points are the raw data for each individual trial, jittered to reduce point overlap

Males made significantly more territorial intrusion calls than did females during both the stimulus and post-trial time periods (Fig. 5; b = 5.087, z = 3.106, *P* = 0.002).

**Fig. 5.**
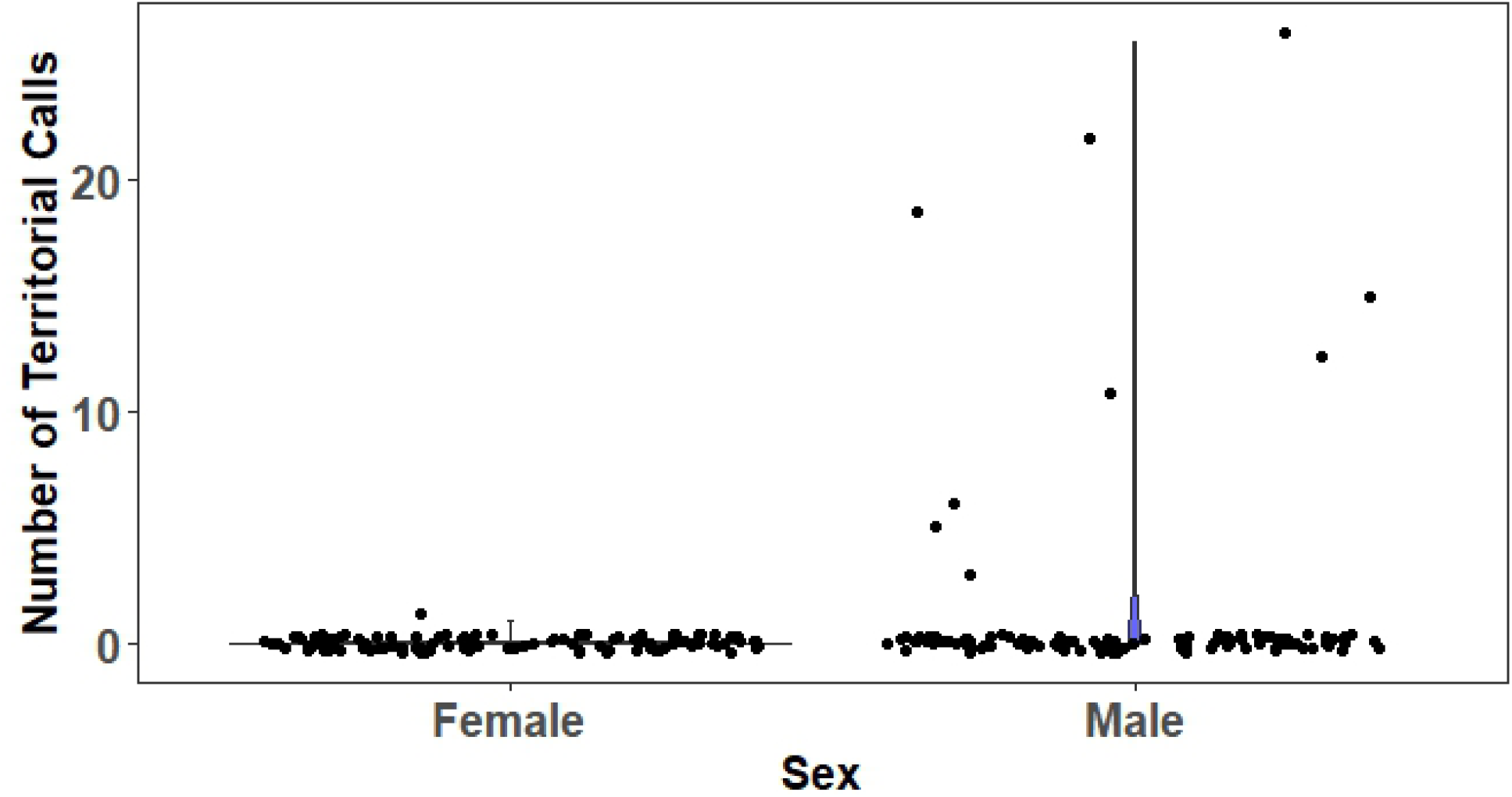
Violin plots and raw data showing the distribution of the number of territorial intrusion calls males and females made during the stimulus and post-trial period after each type of stimulus. The points are the raw data for each individual trial, jittered to reduce point overlap

## Discussion

Species raised under human care have the potential to lose some of their survival-relevant behaviors (McPhee and Carlstead 2010; Shier 2016), including responses to species-specific vocalizations (Corfield et al. 2008; Digby et al. 2013; Tanimoto et al. 2017b). Conservation breeding programs or managed care may also change selective forces operating across generations, or alter developmental processes operating during individuals’ lifetimes, creating new cultural vocal variants and response patterns (Lewis et al. 2021). Disrupting the normal signaler-receiver interplay may result in a breakdown in the communication system, with fitness consequences challenging conservation goals. To address these possibilities, we tested whether pairs of ‘alalā have maintained responses to survival-relevant classes of vocalizations. Although the conservation breeding history of the species may have reduced some of their responses, we still saw encouraging signs that ‘alalā are still able to distinguish between natural call types, and to demonstrate responses that may be linked to increased survival in the wild. Specifically, we found that ‘alalā were more likely to approach in response to alarm calls, indicating they may have been investigating a potential threat or seeking to coordinate a defensive response with conspecifics. Also, we found that males in particular were more likely than females to respond to territorial intrusion calls with territorial intrusion calls of their own, suggesting they were willing to defend their breeding aviary. Finally, our results illustrate that there is some individual consistency in the amount of calling over all treatments, but that individual characteristics (e.g. personality or specific rearing history) do not fully explain the variation in call response to the different treatments. A note of caution in interpreting these results is warranted due to the absence of any quantitative and limited qualitative observations of ‘alalā vocal behavior before they went extinct in the wild, so we have few reference points other than comparisons with and generalizations from other species.

As corvids, flexible learning is expected throughout ‘alalās’ lives (Emery and Clayton 2004), but vocalizations often require exposure during times of parental care or sensitive periods, which may be limited under human care (Corfield et al. 2008; Digby et al. 2013; Tanimoto et al. 2017b). Most ‘alalā in this study were puppet-reared by human caretakers (Greggor et al. 2018), removing any opportunities for learning vocal behavior from parents. Of known call types, alarm calls are more likely to be preserved since they potentially share some innate characteristics that make them harder for predators to locate (Maynard et al. 2003). Additionally, animals including corvids will often make alarm calls to elicit assistance from conspecifics in investigating or confronting a predator (Curio et al. 1978; Coomes et al. 2019). In the case of ‘alalā, we found that birds were more likely to approach alarm call playbacks than other auditory stimuli. However, they were not more likely to make alarm calls in response to alarm playbacks. This lack of vocal response is consistent with what was reported in some pilot work on ‘alalā antipredator behavior (Greggor et al. 2021). This suggests that alarm calls may function to alert birds to the need to gather additional information, or that birds may not respond to alarm calls with their own calls in order to avoid drawing the attention of the threat to themselves. It is also possible that because we presented alarm calls without other stimuli that could indicate danger, they investigated but did not respond with their own alarm calls when they saw no clear threat. Further testing of alarm calls with and without a dangerous context would be necessary to tease this apart. However, ‘alalā’s approach to the source of the playback alone may still have had a social function. The fact that birds approached the alarm calls (and not just conspecific playbacks in general) suggests that the calls may still function to elicit social assistance: approach of the caller may precede mobbing or other group antipredator defense. Mobbing-like behavior has been anecdotally reported in ‘alalā (Greggor et al. 2021); this study provides some empirical evidence that the birds can respond appropriately to alarm calls in the absence of other signs of danger, and thus that they have not become completely desensitized to them.

Territorial calls are also an important part of vocal communication, including in the historical vocal repertoire of ‘alalā (Banko et al. 2002; Tanimoto et al. 2017a). We played two types of territorial calls, a neutral territorial maintenance call and an aggressive territorial intrusion call. ‘Alalā did not significantly differ in their response to the territorial maintenance call from other call types. However, we found that ‘alalā responded to a simulated territorial intrusion with territorial intrusion calls significantly more than they did to any other playback stimuli. A vocal territorial response is the natural reaction we would expect to a territorial challenge (Maynard et al. 2003; Bradbury and Vehrencamp 2011), considering that we presented the birds with this stimulus in their home aviaries. We did not test any of the birds in a new location outside of their artificial “territory” so we do not know if their responses are flexible and adaptive to context, i.e. defending only their occupied territory. However, given that ‘alalā exhibit fewer territorial calls than their wild counterparts (Tanimoto et al. 2017b), we find it encouraging that ‘alalā displayed this behavioral response, suggesting some normal signaler-receiver exchange. Additionally, we found that males were more likely to make territorial intrusion calls, which mirrors patterns noted previously in the wild (Banko et al. 2002). As a sexually dimorphic species with respect to size, males are the larger and more aggressive sex, and thus their greater role in territorial defense is expected (Archer 1988).

While we found encouraging signs that some birds exhibited natural responses to our playbacks in the absence of any other context or stimuli, many of the birds did not respond vocally when faced with any type of conspecific call. Given that the birds are living in social densities that are far higher than observed in the wild (Flanagan et al. 2020), the high levels of individual variation we found suggest that some ‘alalā may have become desensitized to the calls of conspecifics vocalizations, perhaps a result of repeated exposure without consequence, or because salient context was missing from our stimulus presentations. While ‘alalā in the conservation breeding facilities may not necessarily be less vocal than wild birds (Tanimoto et al. 2017b), the lack of vocal behavior we saw in many individuals in response to social cues suggests a potential decoupling of the meaning or relevance of conspecific calls. Alternatively, perhaps there is additional context we did not adequately capture in our recordings (e.g. individual caller ID, social status, etc.), that could have differentially influenced some individuals more than others. Corvids are known to respond to individual qualities of callers (Boeckle et al. 2012), such as their dominance status (Massen et al. 2014) and membership in the breeding colony (Woods et al. 2018), with call signatures that may also help with distinguishing sex (Mates et al. 2015). Additionally, since calls were recorded opportunistically, there may be subtle differences to the different calls (for example, the particular stimulus causing the alarm calls) that we are unaware of. Therefore, we may have inadvertently broadcast information beyond the content of the calls. However, we saw no differences in the number of calls birds made in response to the three different call sets, suggesting that the identity of the caller was not a major cause of variation. A final explanation for variation in response to calls could be that by using recordings of birds in the conservation breeding centers, which was necessary for controlling context and individual factors such as sex and identity, the calls themselves may no longer retain the same information as wild calls would have. Examinations of historical recordings show that similar call types to the ones we broadcast were used by the last wild birds (Tanimoto et al. 2017a). Although some differences likely exist (Tanimoto et al. 2017b), the call types we chose were similar to those produced by wild birds, and the fact that responses were largely consistent with our predictions suggest that the calls used have retained their function. Regardless of the cause of the low responses of many individuals, there were still other birds that clearly demonstrated survival-relevant responses.

To conclude, although we found no evidence of widespread erosion of vocal communication behavior under human care, the individual differences that we saw in how birds responded to the playbacks implies that not all individuals are equally as well equipped for release into the wild, especially if these call responses result in reduced predation risk or increased territory defense in the wild. Many factors go into determining fitness for release, and these results suggest that we may need to consider whether birds demonstrate adequate responses to conspecific calls as a criterion for release. Additionally, future research could investigate how likely birds are to regain survival-relevant responses to vocalizations if they are exposed to training that encourages associations between conspecific calls and relevant responses. Given the critical conservation status of the species, any technique that could limit the impact of the conservation breeding environment on survival-relevant behavior warrants further evaluation.

## Acknowledgments

We greatly appreciate the wildlife care specialists at KBCC and MBCC who accommodated this research and thank Patrick Hart for lending us recording equipment. We would also like to thank members of the Catenazzi and Cox labs for their feedback on an earlier draft and to Ramona Rauber for helpful discussion about ‘alalā call types. We would like to thank Jeffrey Podos in his role as an editor and two anonymous reviewers for their feedback.

## Declarations

### Availability of data and material

Data and code are available on the OSF repository at: https://osf.io/pu5fb/. Pre-print available at: https://www.biorxiv.org/content/biorxiv/early/2021/05/25/2021.05.24.445466.full.pdf

### Code availability

Data and code are available on the OSF repository at: https://osf.io/pu5fb/.

### Ethics approval

This work using animal subjects was approved by San Diego Zoo Wildlife Alliance’s IACUC committee (No. 16-009). All applicable international, national, and/or institutional guidelines for the use of animals were followed. Several permits were issued for the conservation breeding of ‘alala (USFWS Native Endangered Species Recovery Permit TE060179-5, State of Hawaii Protected Wildlife Permit WL19-16).

